# Beta oscillations support somatosensory temporal integration for body ownership

**DOI:** 10.64898/2026.02.03.703611

**Authors:** Mariano D’Angelo, Renzo C. Lanfranco, H. Henrik Ehrsson

## Abstract

The sense of body ownership, the perceptual experience of what belongs to one’s body, relies on the integration of self-related signals across multiple sensory modalities. Alpha oscillations modulate the temporal resolution of visuotactile integration for body ownership by constraining how visual and tactile signals from the body are temporally combined or segregated. However, body ownership can be supported and modulated in the absence of vision, for example during self-touch, raising the question of whether the same alpha-based temporal mechanism supports purely somatosensory integration. Here, we show that individual beta frequency (IBF), rather than alpha, predicts the temporal integration of tactile signals between the two hands in a somatic rubber hand illusion self-touch paradigm. IBF predicted the temporal window within which two tactile stimuli were judged as simultaneous and within which illusory self-touch body ownership was experienced. Moreover, while individual alpha frequency did not contribute to this purely somatosensory integration, IBF also did not predict the temporal resolution of visuotactile integration when visual body-related inputs were available. Together, these findings suggest a double dissociation in which frequency-specific oscillatory processes reflect the temporal integration of bodily signals in a modality-dependent manner: beta frequency emerges as the key correlate of somatosensory integration during self-touch, whereas alpha frequency governs visuotactile integration when vision is available.

## Introduction

The brain reduces the complexity of the sensory environment by temporally binding inputs across time and sensory modalities, thereby forming coherent perceptual objects and events and generating efficient representations of the external world^1^. This process, known as multisensory integration, is tightly linked to the temporal resolution of the sensory systems. Each modality has its own intrinsic temporal precision, and the brain must determine whether inputs from different channels, such as touch, vision and proprioception, are causally related or independent^2^. To solve this problem, the brain evaluates the likelihood that multiple signals originate from a common source, primarily based on their temporal and spatial proximity^3,4^. When the temporal discrepancy between stimuli is small, it is more likely that they stem from the same source and should therefore be integrated. In contrast, when the temporal discrepancy is larger, the signals are more likely to originate from different sources and should be segregated. Therefore, temporal resolution of this integrative mechanism determines which signals are bound together across time and modalities to generate the perception of meaningful objects and events^5–7^. Solving this binding problem is essential also for building a coherent representation of one’s own body^8–13^. To generate the perception of a limb or body part as one’s own, commonly referred to as the sense of body ownership^8^, the brain has to temporally integrate stimuli originating from one’s own body, while distinguishing them from those arising in the external environment^12^. Thus, the multisensory integration processes underlying the sense of body ownership also rely on the specific temporal resolution of the integration process^12,13^. It has been shown that the temporal resolution of visuotactile integration in the context of body ownership correlates with the temporal integration of such inputs in the context of external stimuli^9,12,13^, indicating that similar temporal integration principles determine body ownership perception and perceptual of external events and objects.

Due to their intrinsic rhythmic nature, brain oscillations, particularly in the alpha frequency range, have been proposed as a key mechanism for temporally gating sensory information from the external environment^14–23^. Recent work suggests that alpha oscillations play a central role in discretizing continuous visual input into perceptual frames, effectively shaping the temporal resolution of perception^17^. Faster alpha frequencies are associated with higher temporal resolution, enabling more precise perceptual experiences^18–22^. This relationship implies that individuals with faster alpha rhythms accumulate sensory evidence over a greater number of cycles within a given time window, leading to finer temporal discrimination. Alpha oscillations are most predominantly expressed over occipital^14–22^ and posterior parietal cortices ^22–25^, reflecting contributions from cortical and thalamo-cortical networks. These regions play key roles for visual processing and for the integration of visual information with other sensory modalities, including audition and touch. The involvement of alpha frequency in the temporal integration of perceptual information has been demonstrated primarily in the context of visual processing^16–24^ and audio-visual integration^14,15,26^. Alpha oscillations modulate the temporal window within which stimuli are perceived as unified, commonly referred to as the temporal binding window (TBW). Building on this, a recent study has shown that alpha frequency also plays a role in integrating body-related visual information with somatosensory signals, thereby contributing to visuotactile simultaneity perception and body ownership perception^12^. Together, this body of work raises a fundamental question in perceptual neuroscience: does individual alpha frequency provide a general, amodal constraint on the temporal integration of sensory information, or do different sensory modalities rely on distinct oscillatory frequency bands to support temporal integration?^17^

This question is particularly important for understanding how a unified sense of the body is built from somatosensory inputs alone. Indeed, body ownership is not solely dependent on visual input but can also emerge through the integration of multiple somatosensory signals, such as touch and proprioception, and tactile inputs from two body parts during self-touch^27^. Self-touch is a particularly fundamental cue for bodily self-perception, as it provides highly reliable, temporally and spatially coupled signals originating from the body itself^30–33^. Developmentally, self-touch is among the earliest forms of sensory experience through which the nervous system distinguishes self-generated from external sensory events^34–38^. Thus, while vision often play an important role, a coherent sense of bodily self can be constructed even in absence of vision based on inputs from other modalities^30,39–42^. In such non-visual contexts, the brain relies on temporal and spatial congruency between tactile, proprioceptive, other somatosensory inputs. However, it remains unclear whether the temporal integration of non-visual bodily information, such as touch and proprioception, is supported by the same alpha-based temporal integration mechanism^12^, or whether other frequency bands, such as beta oscillations, which are strongly implicated in somatosensory and motor processing^27–29^, play a more central role. Previous research on brain oscillations and perceptual sampling has focused on contexts in which visual input is available^17^, offering limited insight into how cortical oscillations support multisensory integration when non-visual sensory modalities become more relevant. From this perspective, paradigms based on temporally structured self-touch provide a particularly informative test case for this broader theoretical question, as they isolate somatosensory temporal integration in the absence of vision and allow assessment of whether alpha-based temporal constraints generalize beyond visually related perception.

We hypothesized that beta frequency oscillations support the temporal integration of tactile signals in generating the sense of body ownership in the absence of vision, given that somatosensory processing is typically governed by oscillatory activity in the beta band^27–29^. To test this hypothesis, we developed a robot-assisted psychophysical task designed to measure the temporal integration of bimanual tactile information contributing to body ownership without visual input. Specifically, we implemented a psychophysical version of the somatic rubber hand illusion, which is a well-established paradigm for investigating how tactile and proprioceptive information contribute to the experience of body ownership in the context of self-touch^30,39–42^. In the classical version of the somatic rubber hand illusion, the experimenter guides the blindfolded participants’ left index finger to touch a right rubber hand, while simultaneously delivering tactile stimulation to the participant’s real right hand at the corresponding location. After a brief period of repeated stimulation, most participants report the illusory sensation of directly touching their own right hand with their left index finger, despite physically contacting a rubber hand. Crucially, this illusion breaks down when the movement of the left index finger and its touch feedback are temporally asynchronous with the tactile feedback from the right hand, suggesting that the experience of body ownership in a non-visual context depends on the temporal congruency between tactile signals from the two hands^30^. In our robotically-controlled psychophysical adaptation of this paradigm, we precisely manipulate the timing of tactile events delivered to the rubber hand and the participant’s real hand. We systematically introduced temporal delays or advances of up to 400 ms and asked participants to judge whether they experienced the sensation of touching their own hand or not. To complement the body ownership paradigm, we also included a bimanual tactile simultaneity judgment task as a reference measure of temporal integration in the somatosensory system. This task provides a well-established, assumption-free index of bimanual tactile temporal processing, allowing us to interpret ownership-related effects in relation to basic perceptual integration mechanisms. In this task, participants judged whether the touch they felt from their real left index and the touch they felt from their right hand occurred synchronously or asynchronously. By parametrically modulating the temporal delays (in both tasks), we were able to estimate, for each participant, the level of asynchrony at which simultaneity was perceived, and the illusion of body ownership emerged. To this aim, we computed the TBW for body ownership and simultaneity perceptions and assessed their perceptual sensitivity using a signal detection theory (SDT) approach.

In Experiment 1, we behaviourally investigated temporal resolution of the bimanual integration of tactile information underlying body ownership in the illusory self-touch paradigm and tested whether this resolution correlates with the temporal precision of bimanual tactile simultaneity perception. A positive correlation would indicate a shared temporal tactile integration processes supporting both body ownership and tactile simultaneity perception, consistent with our overarching hypothesis of a common oscillatory frequency mechanism. In Experiment 2, we tested this hypothesis using EEG comparing behavioural measures with oscillatory frequency estimates obtained at rest and during the body ownership and simultaneity tasks, assessing whether alpha- or beta-band frequencies are preferentially associated with the temporal resolution of tactile integration in the two tasks. In Experiment 3 we tested whether the hypothesised beta-frequency relationship is specific to tactile integration in the absence of vision by examining whether beta frequency predicts the temporal resolution of visuotactile simultaneity perception and visuotactile body ownership in a classic visual rubber hand illusion paradigm (based on data from^12^). We hypothesized that this relationship would not extend to visuotactile integration. Collectively, these experiments reveal a modality-specific dissociation in the oscillatory mechanisms supporting bodily self-perception: beta-band frequency selectively governs somatosensory temporal integration during self-touch in the absence of vision, whereas alpha-band frequency supports visuotactile integration when visual information is available. More broadly, these findings suggest that the temporal resolution of bodily multisensory integration is not governed by a single, amodal oscillatory clock, but instead depends on frequency-specific mechanisms associated with the sensory modality through which bodily information is available.

## Results

### Experiment 1: correlation between body ownership and tactile simultaneity TBWs

In Experiment 1, we developed a novel psychophysical paradigm to assess participants’ ability to detect body ownership based solely on tactile and proprioceptive cues, in the absence of visual input. This experiment had two primary aims. First, we sought to determine whether the task could reliably measure the temporal resolution of bimanual tactile integration underlying body ownership by fitting psychometric functions and computing perceptual sensitivity. Second, we investigated whether the temporal resolution associated with body ownership correlated with that of bimanual tactile simultaneity perception. Such a correlation would suggest shared temporal integration processes supporting both the detection of simultaneity and the experience of body ownership.

To these aims, participants performed two psychophysical detection tasks: *(i)* a body ownership judgment task to evaluate the temporal precision of tactile integration underlying body ownership in the absence of vision, and *(ii)* a bimanual tactile simultaneity judgment task to assess participants’ temporal resolution for detecting synchrony between tactile events delivered to the two hands. In both tasks, participants’ left index finger was connected to a robotic arm that guided the finger to touch a rubber hand (**Fig. 1a**). Simultaneously, a second robotic device delivered taps on the participants’ real right hand using a fake finger mounted on the robotic arm. When these tactile events are delivered synchronously, this setup typically elicits the somatic rubber hand illusion^30^, in which participants experience the sensation of directly touching their own hand despite physically interacting with a rubber hand.

**Fig. 1.**
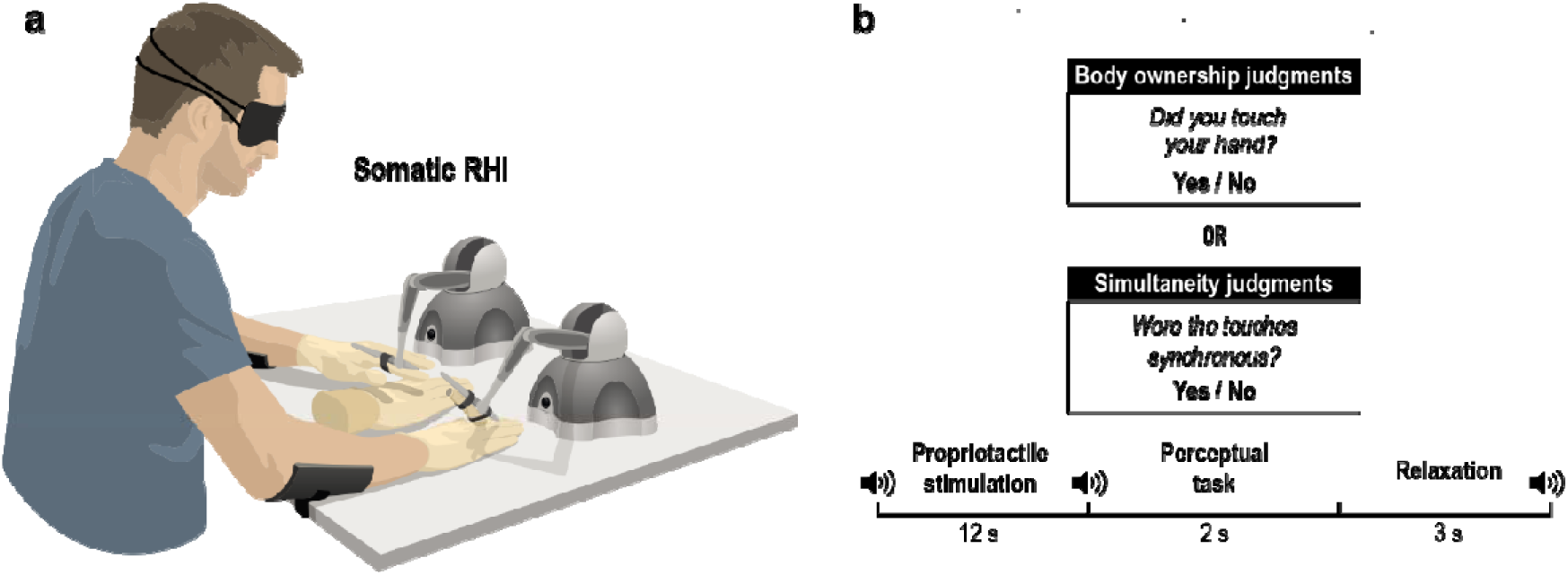
Experimental setup and procedure of Experiment 1. **a,** Experimental setup for the body ownership judgment and simultaneity judgments based on the somatic rubber hand illusion. **b**, Procedure for the body ownership and simultaneity judgment tasks. Participants were blindfolded, with their left index finger connected to a robotic arm that guided the finger to touch a rubber hand. A second robotic device delivered taps on the participants’ real right hand using a fake finger mounted on the robotic arm. Both the rubber hand and the real hand were touched six times over 12-s periods, either synchronously or with the rubber hand being touched slightly earlier or later. The degree of asynchrony was systematically manipulated between 0, ±80 ms, ±160 ms, ±240 ms, and ±400 ms. After each bimanual somatosensory stimulation period, participants indicated either whether the rubber hand felt like their own hand (body ownership judgement task), or whether the touches were synchronized (simultaneity judgement task), using a yes/no forced-choice response. The trials ended with a short relaxation period before the next trial started. The participants were alerted to the different phases of the task with auditory cues (sound symbols).

In our setup, taps on the rubber hand were either synchronized with taps on the participant’s real hand (synchronous condition) or delayed/advanced at four specific asynchronies up to 400 ms. Each trial consisted of six consecutive taps delivered over a 12-s period, a duration sufficiently long to elicit the rubber hand illusion^30,43^. The body ownership judgment and simultaneity judgment tasks were administered in separate blocks of trials. In the body ownership judgment task, in each trial, when the robots stopped, participants were instructed to verbally report whether the rubber hand felt like their own hand by saying “yes” (“I felt like I was touching my right hand with my left index finger”) or “no” (“I did not feel like I was touching my right hand”). In the simultaneity judgment task, when the robots stopped, participants reported whether the touches that they gave on the rubber hand and touches they received on their real hand were synchronous or not by saying “yes” or “no” (**Fig 1b**).

We first confirmed that both tasks exhibited the expected psychometric dependence on temporal asynchrony, providing a basic validation of the psychophysical somatic rubber hand illusion paradigm. Body ownership judgments varied as a function of the delays between the tactile signals delivered to the rubber hand and the participant’s real hand (F_8,192_ = 61.507, p <.001; η^2^_p_= .719; **Fig. 2a**). Similarly, simultaneity reports also varied as a function of temporal delay as expected (F_8,192_ = 75.323, p <.001; η^2^_p_ = .758; **Fig.2b**).

**Fig. 2.**
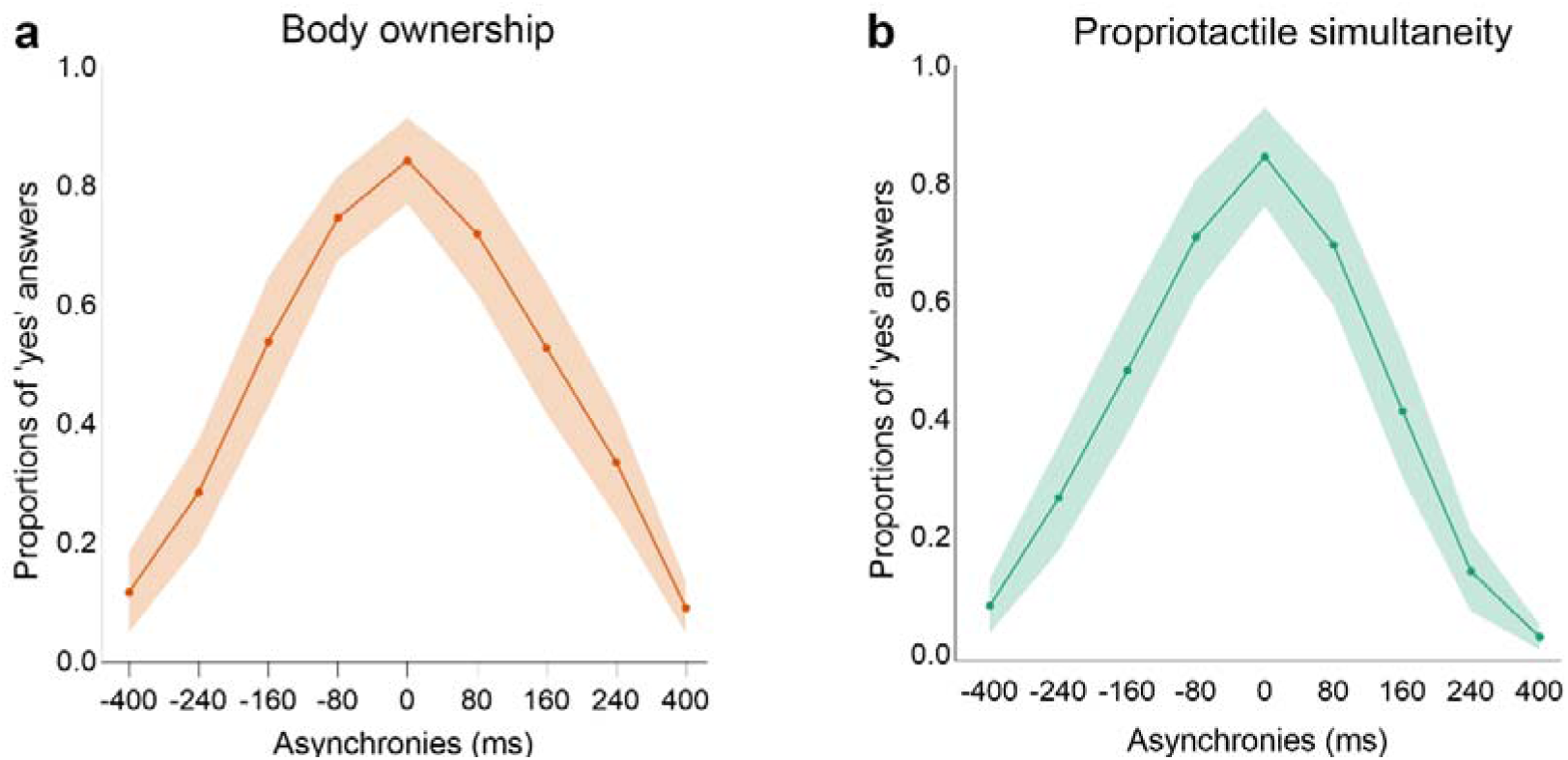
**a,** Proportions of body ownership and **b**, bimanual tactile simultaneity reports as a function of temporal asynchrony. Shaded regions indicate 95% confidence interval. In both tasks, judgments peaked when the two touches were close in time and decreased steadily as larger temporal delays were introduced.

Crucially, in both tasks, we computed the TBW of both body ownership and simultaneity judgments tasks as the standard deviation of the Gaussian curve fitted to the proportion of “yes” responses across the asynchronies^12^. The gaussian model provided a very good fit for both judgment tasks. Importantly, we found a significant correlation between the TBWs derived from the body ownership and tactile simultaneity tasks (Pearson’s r = .682; p < .001; Spearman’s rho = .631; p < .001; **Fig. 3a**), indicating that participants who tolerated larger asynchronies when judging body ownership also tolerated larger asynchronies when judging simultaneity. In addition, body ownership TBW was significantly broader than that for simultaneity judgements (t = 2.286; p = .031), consistent with the fact that body ownership relies on additional contextual multisensory information beyond tactile synchrony^12^, in particular the spatial congruence between the rubber hand and the proprioceptively sensed real hand^47,48^.

**Fig. 3.**
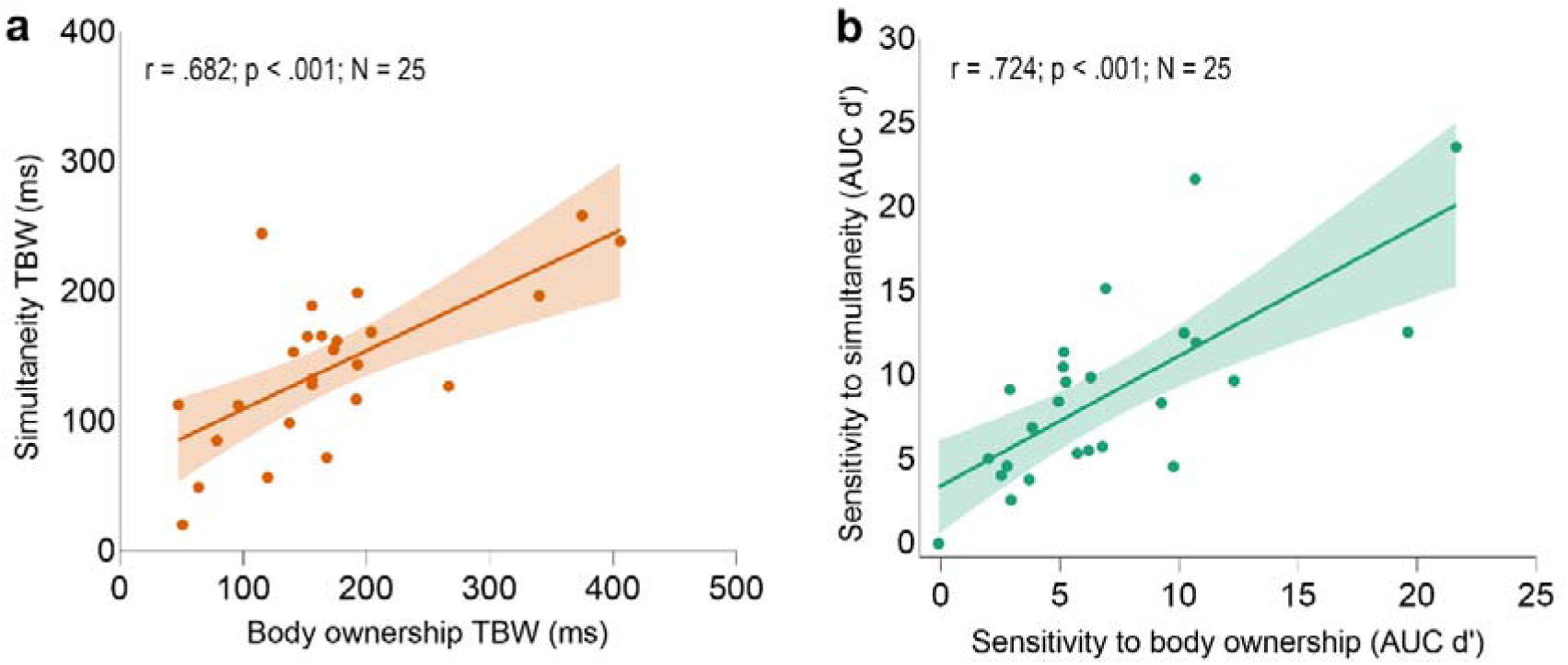
Behavioral correlations in Experiment 1. **a.** Behavioral correlation between body ownership temporal binding window (TBW) and bimanual tactile simultaneity TBW (Pearson’s r =.682). **b**, Behavioral correlation between body ownership sensitivity and bimanual tactile simultaneity sensitivity. In both graphs, the solid line represents the best-fitting regression (Pearson’s r = .724). The shaded region reflects the 95% confidence interval. The AUC denotes the area under the curve of the d’ score measured for each asynchrony.

### Experiment 1: correlation between body ownership and bimanual tactile simultaneity sensitivities

In addition to the TBW, we computed sensitivity (d’) for both the body ownership and simultaneity judgment tasks at each level of asynchrony. Sensitivity was calculated as the difference between the distribution of the probability of hits (i.e., reporting “yes” in the synchronous condition, when no bimanual tactile asynchrony was present) and the distribution of the probability of false alarms (i.e., reporting “yes” in asynchronous conditions, when an asynchrony was introduced) in standardized units^43,44^. This SDT-based analysis allowed us to quantify participants’ sensitivity to simultaneity and to body ownership while controlling for potential bias. As expected, both sensitivity to body ownership (F_3,72_ = 101.293, p <.001; η^2^_p_= .808) and sensitivity to simultaneity increased with greater asynchronies (F_3,72_ = 141.951, p <.001; η^2^_p_ = .855). Crucially, sensitivity to simultaneity judgment tasks significantly correlated with sensitivity to body ownership (Pearson’s r = .724; p < .001; Spearman’s rho = .719; p < .001; **Fig. 3b**). Moreover, sensitivity to simultaneity perception was higher than sensitivity to body ownership (t = 2.247, p = .034), in line with the TBW findings above.

Overall, Experiment 1 shows that the temporal resolution of bimanual tactile simultaneity perception closely related to the temporal resolution of somatosensensory integration underlying body ownership. In Experiment 2, we tested the hypothesis that individual beta frequency (IBF) constitutes an electrophysiological mechanism underlying this behavioral relationship.

### Experiment 2: correlation between parietal IBF and TBWs

In Experiment 2, we tested whether individual beta frequency (IBF) predicts individual differences in temporal integration underlying body ownership and tactile simultaneity perception, and whether these effects differ from those associated with individual alpha frequency (IAF). Prior to the analysis, we defined two regions of interest (ROIs) based on theoretical and anatomical considerations. First, we focused our analysis on a parietal ROI that included the primary somatosensory cortex (S1) and the posterior parietal cortex (PPC) of both hemispheres. Beta oscillations are reliably elicited by tactile stimulation in the somatosensory cortex^45,46^ and the posterior parietal cortex plays a key role in integrating tactile signals from the upper limbs and updating body representation^30,50,51^. Importantly, both alpha and beta frequency peaks can be robustly recorded over the PPC, which exhibits prominent alpha-band activity during multisensory^15,26^ and visuotactile processing^12,25^, and beta-band responses during somatosensory integration and proprioceptive updating^45,52,53^. IAF and IBF were recorded during both resting-state condition and during the execution of the body ownership and simultaneity judgment tasks (**Fig. 4ab**), to test whether the same frequency–behavior relationships hold across resting and task states. We applied the FOOOF/Specparam algorithm to estimate peaks in the alpha and beta frequency bands, controlling for the aperiodic component of the power spectrum^54,55^. This procedure separates true rhythmic activity from the broadband ‘1/f’ background signal, allowing reliable identification of genuine oscillatory peaks. FOOOF successfully identified both IAF and IBF for most of the electrodes within the selected ROIs for all participants (see Methods section).

**Fig. 4.**
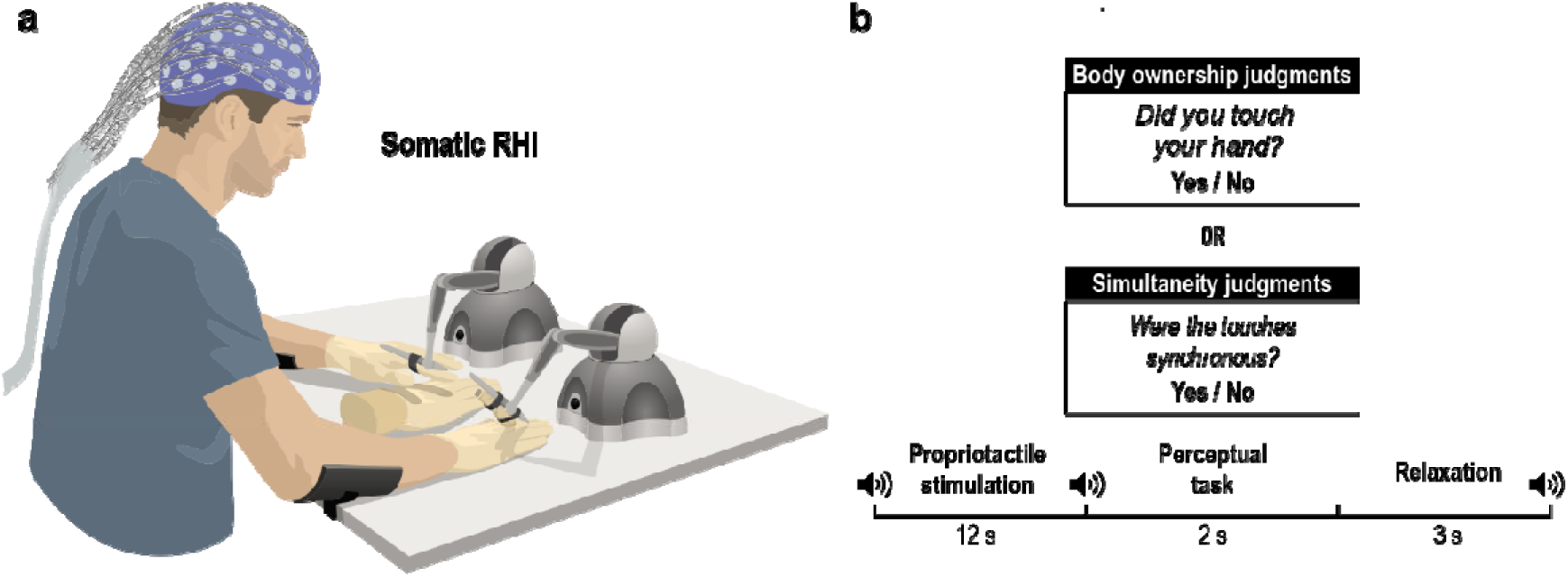
Experimental setup and procedure in Experiment 2. **a,** setup for the EEG Experiment. **b**, procedure for body ownership and simultaneity judgments. Participants kept their eyes closed with their left index finger connected to a robotic arm that guided the finger to touch a rubber hand. A second robotic device delivered taps on the participants’ real right hand using a fake finger mounted on the robotic arm. Both the rubber hand and the real hand were touched six times over 12-s periods, either synchronously or with the rubber hand being touched slightly earlier or later. The degree of asynchrony was systematically manipulated between 0, ±80 ms, ±160 ms, ±240 ms, and ±400 ms.

As predicted, we observed significant negative correlations between parietal resting-state IBF and the body ownership TBW (Pearson’s r = -.484; p =.003; Spearman’s rho = -.464; p = 004; N = 36; **Fig. 5a**), as well as between resting-state IBF and the simultaneity TBW (Pearson’s r = -.478; p = .003; N = 36; Spearmans’ rho = -.482, p = .003). Participants with slower resting-state IBF exhibited broader TBWs for both body ownership and simultaneity judgments. When examining task-related IBF, we again found a significant correlation with body ownership TBW (Pearson’s r = -.529; p = .001; Spearman’s rho = -.441; p = .007; N = 36; **Fig. 5c**). However, the correlation between task-related IBF and tactile simultaneity TBW did not reach significance (Pearson’s r = -.112; p = .517; Spearman’s rho = -.224; p = .188).

**Fig. 5.**
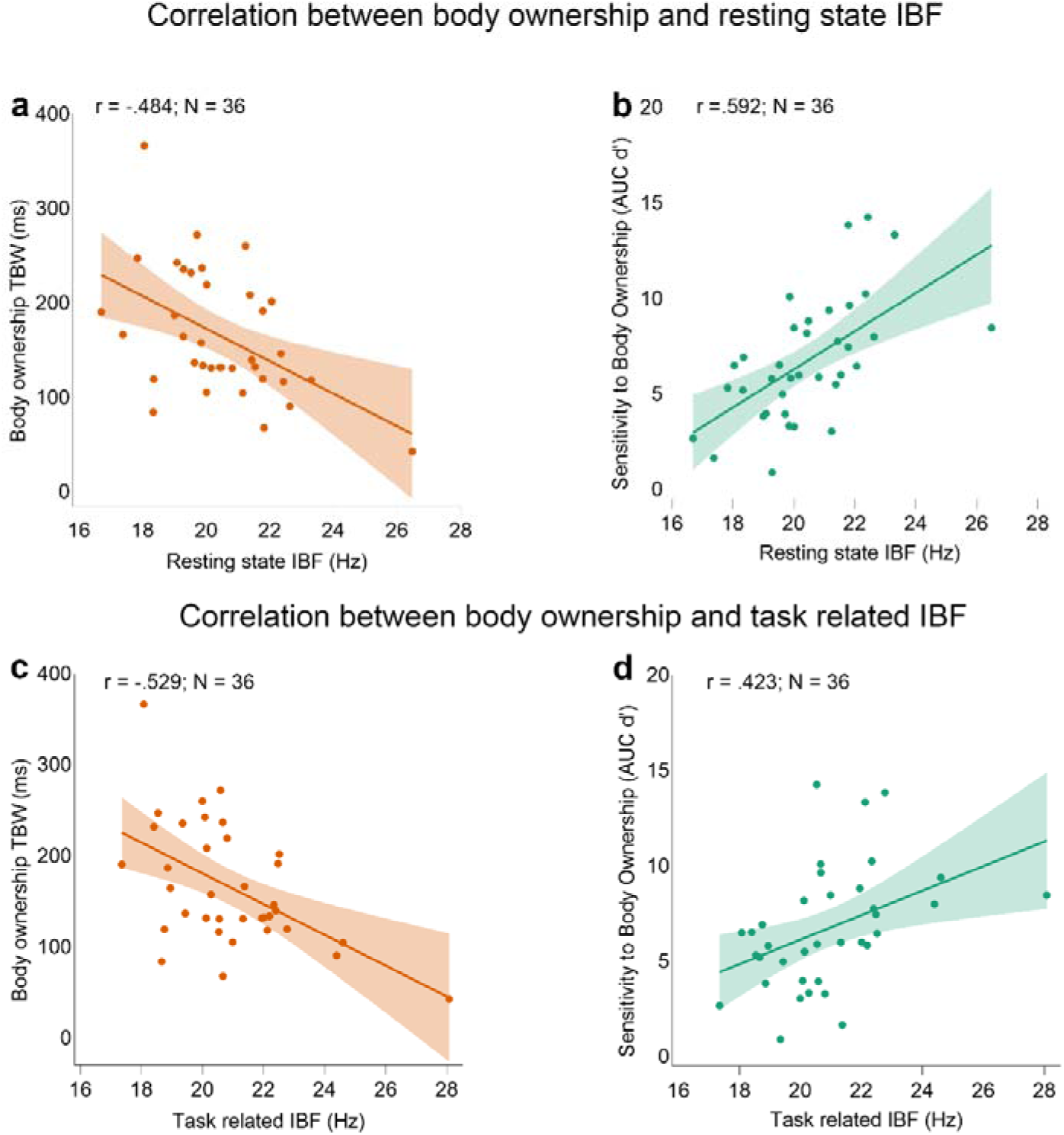
Correlations between individual parietal beta frequency (IBF) and body ownership judgments. **a.** Significant correlation between resting state IBF and body ownership temporal binding window (TBW; Pearson’s r = -.484). **b**. Significant correlation between resting state IBF and body ownership sensitivity (Pearson’s r = .592). **c**. Significant correlation between task related IBF and body ownership TBW (Pearson’s r = .529) **d.** Correlation between task related IBF and body ownership sensitivity (Pearson’s r = .423). In both graphs, the solid line represents the best-fitting regression. The shaded region reflects the 95% confidence interval. The AUC denotes the area under the curve of the d’ score measured for each asynchrony.

In contrast, no relationship emerged between resting-state IAF and body ownership TBW (Pearson’s r = -254, p = .134; Spearman’s rho = -.124, p = .471), nor between resting-state IAF and the simultaneity TBW (Pearson’s r = -.168, p = .326; Spearman’s rho = -.219, p = .200). Similarly, no relationship emerged between task-related IAF and body ownership TBW (Pearson’s r = -.212, p = .215; Spearman’s rho = -.072, p = .678), nor between related IAF and the simultaneity TBW (Pearson’s r = -.040, p = .815; Spearman’s rho = -.136, p = .428).

### Experiment 2: correlation between parietal IBF and perceptual sensitivities

As in Experiment 1, using SDT analysis, we computed body ownership and simultaneity sensitivity, thereby controlling for potential bias. As predicted, we found a significant correlation between resting-state parietal IBF and body ownership sensitivity (Pearson’s r = .592; p <.001; Spearman’s rho = .656; p < .001; **Fig. 5b**), as well as resting-state IBF and simultaneity sensitivity (Pearson’s r = .431; p = .009; Spearman’s rho = .444; p = .007). Moreover, and also as expected, we found a positive correlation between task-related IBF and body ownership sensitivity (Pearson’s r = .423, p = 010; Spearman’s rho = .515, p = 001; **Fig. 5d**). However, this correlation did not reach the significance for simultaneity sensitivity (Pearson’s r = .321, p = .056; Spearman’s rho = .330, p = .050). No correlation emerged between resting-state IAF and sensitivities to body ownership (Pearson’s r = .172, p = .315; Spearman’s rho = .151, p = .378) or between resting-state IAF and tactile simultaneity (Pearson’s r = .243, p = .153; Spearman’s rho = .227, p = .182). Similarly, no relationship emerged between task-related IAF and body ownership sensitivity (Pearson’s r = .040, p = .816; Spearman’s rho = .084, p = .627), nor between task-related IAF and simultaneity sensitivity (Pearson’s r = .209, p = .222; Spearman’s rho = -.192, p = .261).

### Experiment 2: premotor IBF and IAF analyses

We also examined frontal electrodes corresponding to a ROI that covered the bilateral premotor cortex, a region involved in multisensory integration of bodily related inputs and strongly associated with the experience of body ownership in both the visual and somatic variants of the rubber hand illusion paradigms^30,56–58^. Importantly, our recent study using the classical visuotactile rubber hand illusion showed that premotor IAF correlates with both the TBW and body ownership sensitivity^12^, further motivating the examination of this region in the present somatosensory-only context.

We found significant negative correlations between resting-state premotor IBF and body ownership TBW (Pearson’s r = –.580, p < .001; Spearman’s ρ = –.535, p = .001; N = 36; **Fig. 6a**), as well as between resting-state premotor IBF and the simultaneity TBW (Pearson’s r = –.366, p = .028; Spearman’s ρ = –.439, p = .007; N = 36). When considering task-related IBF, we also found significant negative correlations with both body ownership TBW (Pearson’s r = –.476, p = .003; Spearman’s ρ = –.405, p = .014; N = 36; **Fig. 6c**) and simultaneity TBW (Pearson’s r = –.371, p = .026; Spearman’s ρ = –.588, p < .001). In contrast, the correlation with resting-state premotor IAF did not reach significance with either body ownership TBW (Pearson’s r = -.291, p = .085; Spearman’s rho = -.187, p = .275) or simultaneity TBW (Pearson’s r = -.226, p = .185; Spearman’s rho = -.220, p = .197). Similarly, no relationship emerged between task-related IAF and body ownership TBW (Pearson’s r = -.246, p = .148; Spearman’s rho = -.151, p = .380), nor between task-related IAF and simultaneity TBW (Pearson’s r = -.030, p = .864; Spearman’s rho = -.082, p = .636).

**Fig. 6.**
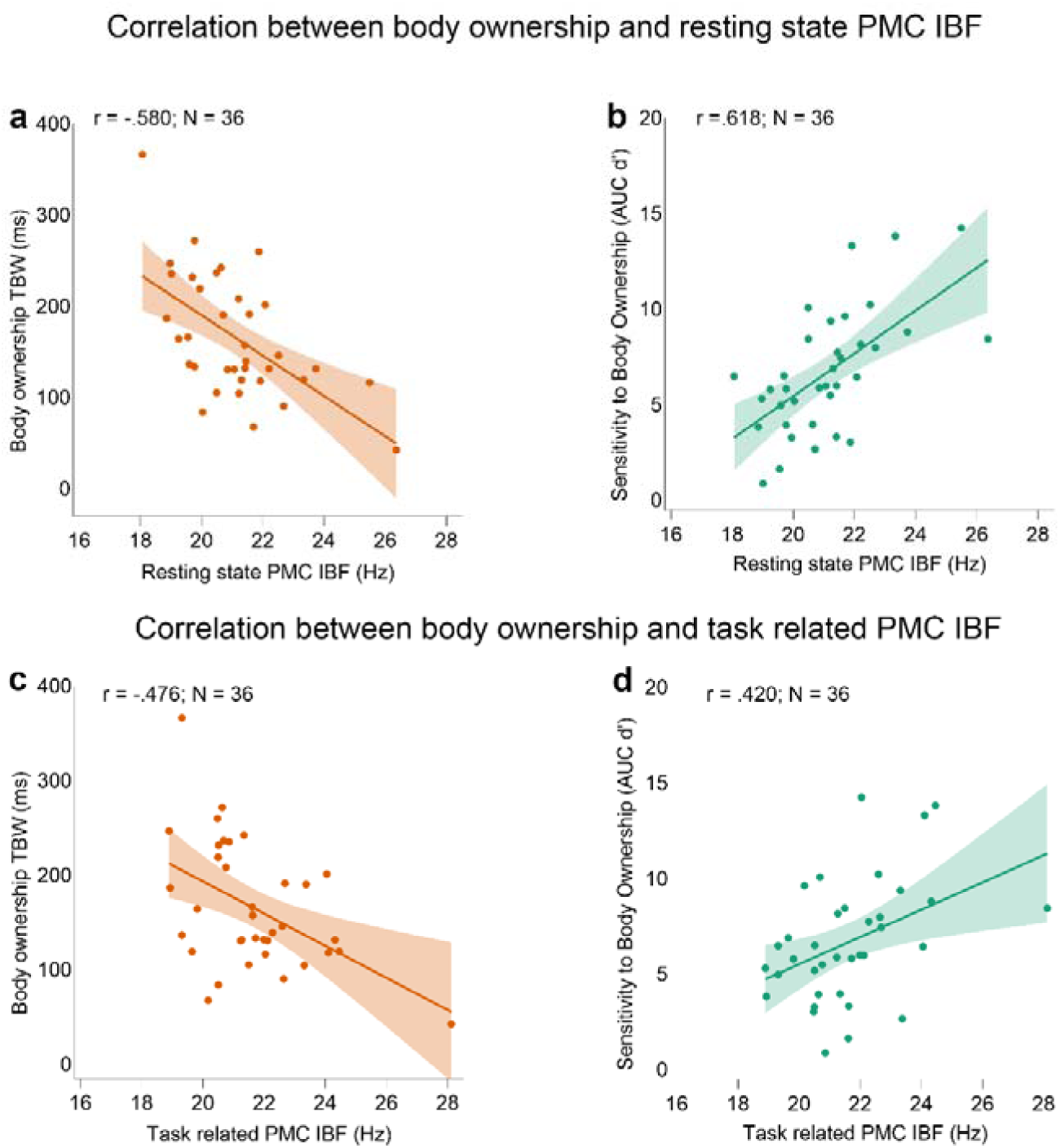
Correlations between individual beta frequency (IBF) from the premotor cortex (PMC) and body ownership judgments. **a.** Significant correlation between resting state IBF and body ownership temporal binding window (TBW; Pearson’s r = .580). **b**. Significant correlation between resting state IBF and body ownership sensitivity (Pearson’s r = .618). **c**. Significant correlation between task related IBF and body ownership TBW (Pearson’s r = .476) **d.** Correlation between task related IBF and body ownership sensitivity (Pearson’s r = 420). In both graphs, the solid line represents the best-fitting regression. The shaded region reflects the 95% confidence interval. The AUC denotes the area under the curve of the d’ score measured for each asynchrony.

Next, we computed body ownership and simultaneity sensitivity to control for bias. We found a significant correlation between resting-state premotor IBF and body ownership sensitivity (Pearson’s r = .618; p <.001; Spearman’s rho = .636; p < .001; **Fig. 6b**). However, the correlation between resting-state IBF and simultaneity sensitivity did not reach significance (Pearson’s r = .299; p = .076; Spearman’s rho = .306; p = .070). When considering task-related premotor IBF, we found a positive correlation with both body ownership sensitivity (Pearson’s r = .420, p = 011; Spearman’s rho = .450, p = 006; **Fig. 6d**), and tactile simultaneity sensitivity (Pearson’s r = .478, p = 003; Spearman’s rho = .463, p = 004). In contrast, no significant correlation emerged between resting-state premotor IAF and sensitivities to body ownership (Pearson’s r = .235, p = .168; Spearman’s rho = .249, p = .143) or tactile simultaneity (Pearson’s r = .313, p = .063; Spearman’s rho = .322., p = .055). Similarly, no relationship emerged between task-related IAF and body ownership sensitivity (Pearson’s r = .016, p = .928; Spearman’s rho = .093, p = .589), nor between task-related IAF and simultaneity sensitivity (Pearson’s r = .265 p = .119; Spearman’s rho = .204, p = .233).

### Experiment 3: no correlation between IBF and temporal integration of visuotactile signals underlying body ownership

Experiment 2 showed that IBF, but not IAF, was associated with the temporal resolution of somatosensory integration underlying body ownership and tactile simultaneity perception in the absence of vision. The absence of a corresponding relationship with IAF is consistent with the hypothesis that the dominant oscillatory mechanisms underlying self-related multisensory integration depend on the specific sensory channels through which the body is perceived. When the body is perceived solely through somatosensory signals, beta frequency appears to be more relevant for shaping the temporal integration of body-related inputs, whereas alpha oscillation frequency is important for the temporal integration of visuotactile bodily signals, as we recently reported^12^. This pattern suggests a functional dissociation between IBF and IAF. In Experiment 3, we further tested this hypothesized dissociation by investigating whether IBF predicts the temporal resolution of visuotactile integration underlying the sense of body ownership in the rubber hand illusion elicited by temporally manipulated visual and tactile inputs.

To this aim, we reanalysed the data from our recent study^12^ to examine IBF in the context of the visual version of the RHI paradigm, which was not assessed in the original analysis. In this study, participants performed a body ownership judgment task and a visuotactile simultaneity judgment task. Specifically, two robotic arms applied six repetitive tactile stimuli to the index finger of a rubber hand in view and to the participant’s hidden real index finger. When the touches on the participant’s hand and the rubber hand were synchronized, most participants report their perception that the rubber hand feels like their own (in the majority of trials). As in the somatic psychophysical paradigm, the taps were delivered either synchronously or with systematic temporal delays, here up to 500 ms. In each trial, when the robots stopped, participants were instructed to verbally report whether the rubber hand felt like their own hand by saying “yes” (“the rubber hand felt like it was my hand”) or “no” (“the rubber hand did not feel like it was my hand”). In the simultaneity judgment task, the visuotactile stimulation was identical to that used in the body ownership task, except that participants reported whether the seen and felt touches were simultaneous. Given that visuotactile body ownership and simultaneity judgments have previously been shown to depend on IAF ^12^, we tested the critical prediction that IBF should not predict temporal integration or perceptual sensitivity in this visuotactile context.

To address this question, we applied the FOOOF/Specparam algorithm to detect IBF over the same parietal ROI as used in Experiment 2 corresponding to bilateral somatosensory and posterior parietal cortices. No significant relationships were found between resting-state IBF and the TBW for body ownership (Pearson’s r = 124, p = .412; Spearman’s rho = .058, p = .701) or for visuotactile simultaneity (Pearson’s r = –.164, p = .277; Spearman’s rho = –.089, p = .554). Likewise, task-related IBF did not correlate with the TBW for body ownership (Pearson’s r = .130, p = .388; Spearman’s rho = .071, p = .638) or TBW for visuotactile simultaneity (Pearson’s r = .002, p = .987; Spearman’s rho = –.136, p = .366). Analyses of perceptual sensitivities yielded consistent results: resting-state IBF did not correlate with sensitivity to body ownership (Pearson’s r = -.017, p = .909; Spearman’s rho =-.091, p = .548) or visuotactile simultaneity (Pearson’s r = -.091, p = .546; Spearman’s rho = -.161, p = .284), and no significant relationships emerged between task-related IBF and sensitivity to body ownership (Pearson’s r = –.250, p = .094; Spearman’s rho = –.166, p = .270) or visuotactile simultaneity (Pearson’s r = –.137, p = .365; Spearman’s rho = .046, p = .759).

Similar negative results were obtained when we analysed IBF from the premotor ROI in the visuotactile RHI data. Resting-state premotor IBF did not correlate with TBW for body ownership (Pearson’s r = –.122, p = .419; Spearman’s rho = –.054, p = .723) or simultaneity judgments (Pearson’s r = –.031 p = .836; Spearman’s rho = –.053, p = .725), nor with sensitivity to body ownership (Pearson’s r = .065, p = .668; Spearman’s rho = –.026, p = .864) or simultaneity (Pearson’s r = .035, p = .816; Spearman’s rho = –.040, p = .793). Similarly, task related premotor IBF did not correlate with TBW for body ownership (Pearson’s r = .133, p = .379; Spearman’s rho = .134, p = .375), nor with its perceptual sensitivity (Pearson’s r = -.243, p = .104; Spearman’s rho = –.154 p = .307). Likewise, no relationship emerged between task-related IBF and the TBW (Pearson’s r = -.090, p = .554; Spearman’s rho = –.094, p = .534) or sensitivity (Pearson’s r = .123, p = .415; Spearman’s rho = .129, p = .394) for simultaneity judgments.

Taken together, these findings indicate that the sensory modality through which one’s body is perceived determines the oscillatory mechanisms supporting the temporal integration of bodily self-related signals. Beta frequency appears to govern tactile integration in the absence of visual information, whereas alpha frequency becomes the key oscillatory rhythm mediating the visuotactile integration when visual and tactile signals are combined or segregated.

## Discussion

In this study, we show that the temporal integration of bimanual tactile signals underlying illusory self-touch body ownership, in the absence of vision, is governed by beta-band oscillatory frequency, and not by alpha frequency. In Experiment 1, we found that the TBW and sensitivity to bimanual tactile simultaneity predicted the TBW and sensitivity of body ownership, suggesting a shared temporal integration mechanism. In Experiment 2, we showed that IBF predicted the temporal integration of bimanual tactile signals underlying body ownership both at rest and during task performance, and that resting-state IBF also predicted tactile simultaneity perception. Importantly, these relationships were specific to beta frequency relative to alpha frequency, as no comparable associations were found for IAF in either parietal or premotor regions. In Experiment 3, we found that IBF did not correlate with TBWs or sensitivities related to visuotactile simultaneity or visuotactile-induced body ownership. Taken together, these findings suggest a modality-specific dissociation in the oscillatory mechanisms supporting body ownership: beta-band frequency governs temporal integration of somatosensory inputs during self-touch, whereas alpha-band frequency supports visuotactile integration^12^.

To generate the sense of body ownership, the brain must selectively integrate signals originating from the body while segregating those arising from external sources^8–12^. Previous studies have shown that the temporal resolution of visuotactile simultaneity correlates with visuotactile integration responsible for the sense of body ownership^12,13^. Experiment 1 extends these findings to the somatic, non-visual, domain by providing the first psychophysical characterization of the somatic rubber hand illusion, demonstrating that illusory self-touch body ownership is systematically governed by bimanual tactile asynchrony. Specifically, we found that body ownership TBW correlated tightly with bimanual tactile simultaneity TBW, and that participants with higher perceptual sensitivity for body ownership exhibited higher sensitivity to bimanual tactile simultaneity. These results provide quantitative psychophysical evidence that body ownership mediated by illusory self-touch in the absence of vision is closely linked to bimanual tactile integration, extending earlier work that relied primarily on categorical synchrony manipulations, questionnaire reports, and indirect measures such as proprioceptive drift^30,41^. Importantly, these findings indicate that body ownership and tactile simultaneity judgments share a common temporal constraint in the somatic domain. This quantitative psychophysical link provides a behavioural foundation for asking how such temporal integration is implemented at the neural level, a question we address in Experiments 2 and 3 by relating individual differences in temporal integration to oscillatory frequency.

Experiment 2 revealed that beta, but not alpha, oscillations support the temporal integration of bimanual tactile information underlying body ownership and simultaneity perception. Beta oscillations are known to originate from the somatosensory cortical system and are integral to a broader sensorimotor network that includes the parietal cortex, the motor cortex, basal ganglia, thalamus, and cerebellum^42–46,58^. These oscillations are closely linked to tactile processing and play a key role in sensorimotor control^60–62^.Touching the skin leads to a desynchronization in beta frequency bands, reflecting a preparation of the somatosensory system for efficient processing of incoming stimuli^45^. Similarly, a reduction of beta power across sensorimotor areas is observed just prior to and during movement execution, indicating a shift from a stable, predictive state to one of increased cortical excitability and readiness for action^63^. A few studies have suggested that beta frequencies, similar to alpha rhythms in the visual domain, might help define discrete perceptual cycles in the somatosensory system, implying that tactile perception may operate in a periodic sampling mode rather than as a continuous process^64–66^. This perspective proposes that the brain does not process tactile input in a continuous stream but instead segments it into temporal windows governed by ongoing beta oscillations, similar to the relationship between alpha frequency rate and visual perception. Our EEG results support this view and suggest that beta frequency contributes to temporal sampling processes that shape the integration of self-related somatosensory signals underlying both body ownership and simultaneity perception during self-touch in the absence of vision.

Our EEG results provide new insights into the role of the PPC and premotor cortex in multisensory bodily integration and body ownership. Beta oscillations recorded over PPC support the integration of tactile information with proprioceptive signals about body position, allowing the brain to construct a coherent and accurate representation of where touch occurs in space^38^. This region is a central hub for multisensory integration^67–69^ and is known to be active in both the visuotactile^10,12,56–58^ and somatic versions^30^ of the rubber hand illusion. Although in both illusions tactile stimuli play a crucial role, the frequency bands responsible for temporal integration vary depending on only somatosensory signals are integrated or also visual signals, according to our findings. In the visuotactile rubber hand illusion, visual input is heavily weighted in the multisensory integration process and vision also provides the primary spatial reference for integrating visual, tactile, and proprioceptive signals, leading to a recalibration of both the perceived location of touch and the felt position of the real hand toward the rubber hand^70,71^. Our previous study linked such visuotactile integration to parietal alpha oscillations^12^. In contrast, the somatic rubber hand illusion relies primarily on the temporal integration of bimanual tactile signals and proprioceptive cues, and participants are blindfolded or have their eyes closed minimizing the role of visual input^30,39–41^. In this illusion, the tactile sensations from the left index finger and the right real hand are experienced as a single self-generated touch event, as if the participant were directly touching their own hand. This unified percept leads to a recalibration of the perceived location of touch on the real hand and an updating of the proprioceptive representation of hand position in space^30,39^. Our results suggest that this form of somatic integration is associated primarily with beta oscillations, a frequency band strongly linked with somatosensory and sensorimotor processing^45–47,52,53^, which may support the temporal integration of tactile inputs from the upper limbs into a coherent multisensory experience of self-touch. Thus, although oscillatory activity in the parietal cortex contributes to both visuotactile and bimanual tactile integration in body ownership, alpha and beta frequencies appear to play distinct roles in these two modes of multisensory bodily integration, with parietal beta contributing specifically to body ownership during self-touch in the absence of vision.

Analysis of frontal electrodes corresponding to the PMC revealed a similar significant relationship between IBF and body ownership and simultaneity TBWs, as well as perceptual sensitivities, in the current somatic RHI paradigm. The PMC is a multisensory area^68, 72,73^ strongly involved in the experience of body ownership^69,70,74^. Importantly, PMC activity has been observed in both the visuotactile^56–58^ and the somatic^30^ versions of the rubber hand illusion, supporting its role in integrating multisensory inputs as well as maintaining sustained self-attribution of the hand^58^ and supporting causal inference of body ownership^57,58^. In our previous study^12^ on the visuotactile RHI, we found that IAF over the PMC correlated with body ownership TBW, but not with visuotactile simultaneity TBWs, in line with a more selective involvement in visuotactile integration in the context of body ownership^12^. In contrast, in the current somatic RHI study we found that PMC IBF predicted both body ownership and simultaneity judgments. This indicates that oscillatory beta activity in the PMC during the somatic RHI support the temporal integration of individual tactile events, shaping both body ownership and simultaneity perception. This observation underscores the role of premotor beta in temporal integration somatosensory signals from the two hands and the significance of this integration of sensing body ownership through self-touch. In both parietal and premotor cortices, beta-band activity, similarly to alpha for vision^75,76^, may set this temporal integration window by rhythmically modulating neuronal excitability, thereby gating when spikes can effectively contribute to tactile stimulus encoding and integration.

A long-standing question in the study of perceptual sampling is whether individual alpha frequency provides a general, amodal mechanism for temporal integration across sensory modalities, or whether different sensory channels rely on distinct oscillatory frequencies^17,65^. The present findings resolve this issue in the context of bodily self-perception by demonstrating a modality-specific dissociation: beta-band frequency governs temporal integration during purely somatosensory self-touch, whereas alpha-band frequency supports visuotactile integration when visual information is available. Previous work, including our own^12^, has established that individual alpha frequency shapes the temporal binding window for visual^18–24^, audiovisual^14,15,26^, and visuotactile perception^25^, including visuotactile body ownership^12^. However, in all of these paradigms, visual input was present and likely dominated temporal integration. In contrast, the somatic rubber hand illusion and the bimanual tactile simultaneity task eliminate visual input entirely, isolating temporal integration within the somatosensory domain. Under these conditions, individual beta frequency, but not alpha frequency, predicted both temporal binding windows and perceptual sensitivity, consistent with evidence that beta oscillations play a central role in somatosensory and sensorimotor processing. This interpretation is further supported by evidence from the tactile version of the sound-induced flash illusion, a cross-modal visual illusion in which the temporal binding window has been shown to correlate with individual beta frequency rather than alpha frequency, consistent with a somatosensory-driven integration process^65^. Together, these results argue against a single, amodal oscillatory clock governing perceptual cycles and instead support a modality-specific account in which temporal integration is shaped by frequency bands tuned to the dominant sensory channel engaged in the task. At a broader level, these results support the view that cortical networks achieve temporal integration through frequency-specific dynamics that are tuned to the sensory pathways and functional networks engaged, rather than through an amodal oscillatory clock, consistent with recent neurophysiological evidence that cortical neurons integrate information within constrained temporal windows^75^.

In summary, our study shows that IBF, but not IAF, predicts the temporal integration of somatosensory signals that contribute to the experience of body ownership driven by self-touch simultaneity cues when vision is absent. In contrast, IBF does not predict the temporal resolution of visuotactile integration, which has instead been linked to individual alpha frequency in previous work^12^. By identifying oscillatory frequency as an electrophysiological mechanism associated with constraints on temporal integration during self-touch perception, our findings highlight self-touch–related somatosensory integration as a fundamental mechanism through which the brain constructs the bodily self. More broadly, these results establish oscillatory frequency as a central dimension for understanding how cortical networks implement temporal integration in a modality-specific manner, rather than through a single amodal timing mechanism.

## Methods

### Experient 1

#### Participants

Thirty-three participants were recruited for the experiment (16 females; ages 18-43 y; mean age: 26.90 years). However, since the ability to experience the somatic rubber hand illusion was required to perform the task and to fit the psychometric functions to the response data, we included in the main experiment only participants (N = 25; 13 females; ages 18-43 y; mean age 27.92 y) that fulfilled the inclusion criterion of being able to experience this variant of the rubber hand illusion (participants’ inclusion test in the Supplementary Material). All participants had no history of neurological or psychiatric disorders and provided written consent to take part in our experiment, which was approved by the Swedish Ethical Review Authority. Participants received a modest financial compensation for their time. We performed a power analysis for sample size estimation using G*Power^78^. Based on previous findings in our lab, we specified a one-tailed correlation test, anticipating a significant positive relationship, a large effect size of 0.5 (Cohen’s d) and a power level of 0.80. The analysis indicated a critical sample size of 21 subjects, which was increased to 25 to identify a correlation between tasks with a power greater than 0.80 and a strong effect size.

#### Experimental setup and procedure

The experiment took place in a quiet, dimly lit room specifically equipped for psychophysical testing. Participants completed two tasks: (i) a body ownership judgment task and (ii) a bimanual tactile simultaneity judgment task. Both tasks involved the use of two 3D Systems Touch haptic devices (https://www.3dsystems.com/haptics-devices/touch). Participants sat comfortably in front of a table covered with a black cloth. Beneath the cloth, a slightly convex and soft surface made of foam and modeling clay was positioned to allow participants to rest their right hand comfortably, palm down, with the fingers naturally curved. They were blindfolded during the task and wore white gloves on both hands. Their left index finger was secured with a velcro strap to a custom-made support attached to the stylus of one haptic device. The finger was held horizontally and guided by the device to perform vertical tapping movements of approximately 2.5 cm in amplitude. Participants tapped on a soft silicone hand covered with a white glove. The distance between the participant’s real right index finger and the corresponding finger of the rubber hand was approximately 9 cm.). A second haptic device held a fake wooden finger, likewise covered with a piece of the same white glove and fitted with a silicone layer beneath the glove at the contact site with the participant’s real hand. This device was used to apply taps to the dorsum of the participant’s right hand, at a soft-tissue location approximately 2 cm proximal to the midpoint between the third and fourth metacarpal bones, to ensure uniform tactile stimulation. Thus, during each trial, one haptic device guided the participant’s tapping movement onto the rubber hand, while the other delivered tactile stimulation to their real hand. A chin rest and two elbow supports ensured that the participant’s head and arms remained stable and relaxed throughout the tasks. Participants also wore earplugs to minimize distraction from any noise produced by the haptic devices. Taps on the rubber hand and the participant’s real hand were either temporally aligned (synchronous condition) or presented with a temporal offset (asynchronous conditions). Asynchronies varied across trials in five levels: ±400 ms, ±240 ms, ±160 ms, ±80 ms, and 0 ms (true synchrony). Negative values indicated that the tap on the rubber hand preceded the tap on the real hand, whereas positive values indicated the opposite. Asynchronies were selected based on a pilot experiment involving 6 participants who performed the tactile simultaneity task.

In each trial, the haptic devices tapped the rubber hand and the participant hand six times each for a total period of 12 s. The interval between consecutive touches was jittered between 1100 and 1300 ms. Then, the robots stopped while the participant heard a tone instructing them to verbally provide their answers. In the body ownership judgment task, they indicated whether they had the sensation to touch their right hand with their left index finger by saying “yes” (“It felt like I was touching my own right hand ”) or “no” (“It did not feel like I was touching my right hand”); in the simultaneity judgment task, participants reported if the tap they gave with the left index and the touch they received on the right hand were simultaneous or not. Five seconds later the end of the stimulation period, a second tone informed the participant that the next trial was about to start, and the next trial started 1 s after this sound cue. Body ownership and simultaneity judgment tasks were administered in three different blocks of trials using the ABAABB counterbalancing order, with the order counterbalanced across participants. Each asynchrony was repeated 5 times in each block, leading to a total of 45 trials per block. In total, each asynchrony was repeated 15 times in both the body ownership and the simultaneity judgments.

#### Data Analysis

To test whether temporal delays between the tactile stimuli modulated body ownership and simultaneity judgments, as expected^9,12^, we run two ANOVAs, one for each task, with temporal delays as a with-subjects factor. For each task, the TBW was then obtained as the standard deviation of the Gaussian curve fitted on the “yes” responses as a function of temporal asynchrony. Therefore, in the simultaneity judgment tasks, the TBW represents the standard deviation of the Gaussian curve fitted to participants’ reports of synchrony. In the body ownership judgment tasks, the TBW was the standard deviation of the Gaussian curve fitted to the trials in which participants reported the feeling of touching their own real hand. We ascertained the presence of a correlation between the two TBWs through Pearson and Spearman correlation analyses. Moreover, we compared the width of body ownership and simultaneity TBWs using a simple paired t-test, as we expected the TBW to be larger for body ownership^12^

In addition to the TBW, we computed d’ scores for each of the asynchronous conditions, collapsing positive and negative asynchronies (≠ 0 ms; noise) with respect to the synchronous condition (= 0 ms; signal). Here, d’ was calculated as d’ = z(H) – z(FA), where z(H) represents the z score of the hit rate, i.e., the probability of correct reactions (i.e., “it felt like I was touching my right hand” in the body ownership judgment task or “the touches were synchronous” in the synchronicity judgment task) in 0-ms trials, and the z(FA) represents the z score of false alarms in asynchronous trials. To avoid zero counts, we applied padding (edge correction) by either adding or subtracting half a trial. We checked whether the sensitivities to body ownership and simultaneity judgment tasks increased with increasing asynchronies, as expected from previous studies^11,12^, with two separate ANOVAs on the d’ scores with asynchronies as a factor. Finally, we computed the area under the curve of the d’ values as a function of the asynchronies (AUC d’) to provide an index of the sensitivity to the simultaneity and body ownership judgment tasks. We ascertained a correlation between body ownership and simultaneity sensitivities by computing Pearson and Spearman correlations on the AUC d’ scores. Additionally, we compared sensitivity to body ownership and sensitivity to simultaneity using a paired sample t test.

### Experiment 2

#### Participants

Forty-five participants were recruited for Experiment 2. However, as for Experiment 1, only participants (N = 36; 18 females; ages 18-39 y; mean age 25.97 y) that fulfilled the inclusion criterion of being able to experience the somatic rubber hand illusion, were included in the study. The power analysis was identical for Experiment 1. However, we decided to increase the sample size to 36 participants to account for the posibility that the FOOOF/Specparam method might fail to detect a clear task-related alpha peak in 10–20% of the participants. All participants had no history of neurological or psychiatric disorders and provided written informed consent to participate in the study. The experiment was approved by the Swedish Ethical Review Authority, and the participants were provided with a modest financial compensation for their participation.

#### Experimental procedure

Experiment 2 was conducted in a dedicated EEG laboratory, a sound-attenuated, windowless room designed for EEG and psychophysical experiments. Before starting the perceptual tasks of the main experiment, we recorded eight minutes of resting-state EEG data: four minutes with eyes closed and four minutes with eyes open. During the resting state period, participants sat on a chair in the same room where the main experiment took place and were instructed to remain still and relaxed. In the eyes-open condition, they were asked to focus on a fixation cross positioned on the wall approximately 75 cm away. In the eyes-closed condition, participants were asked to keep their eyes gently closed while remaining relaxed and refrain from making any movements. Given that the main task was performed with eyes closed, we focused our analyses on the eyes-closed resting-state data. Following the resting-state recording, participants completed the body ownership and simultaneity judgment tasks. The experimental setup and procedure were identical to those used in Experiment 1, including the placement of the rubber and real hands and the use of the robotic arms. Importantly, as in Experiment 1, the interval between consecutive touches was jittered between 1100 and 1300 ms to prevent neural oscillatory entrainment caused by regularly timed stimulation.

#### EEG recording and preprocessing

Both resting-state and task-related EEG data were recorded and digitized at a sampling rate of 1024 Hz using a 128-electrode Biosemi system with an elastic cap, in which electrodes were integrated at sites conforming to the ABC system. All impedance values were kept below 50 kΩ. Scalp electrodes were referenced to A1. Participants were not blindfolded (as in Experiment 1), as this would have interfered with electrode placement; instead, they were instructed to keep their eyes closed during the tactile stimulation, which was continuously monitored by the experimenter throughout the session. The continuous EEG data were resampled to 512 Hz and then filtered, leaving frequencies between 0.3 and 40 Hz, and epoched from -500 ms before to 500 ms after each touch was applied to the participant’s real hand. Electrodes with a variance smaller than -2 or larger than 2 standard deviations of the mean activity of all electrodes were rejected. To identify EEG artifacts, we implemented an independent-component analysis (ICA^79^) on the epoched data. We evaluated each component using the automated machine learning-based ICLabel classifier^80^ and rejected all components that exhibited a 75% or higher probability of being associated with eye blinks, ocular movements, muscle activity, heartbeat, or channel noise. EEG electrodes previously rejected were replaced with the weighted average activity in all remaining electrodes using spherical spline interpolation. On average, 13% [SD = 8] of electrodes were rejected and subsequently interpolated per participant. However, no electrodes were rejected in our two ROIs. Epochs with voltages exceeding ±150 µV were excluded from further analysis; on average, 1.4% [1.6] of trials were excluded per participant. Finally, the EEG signal was re-referenced to the average across all electrodes. Preprocessing was performed using custom-made MATLAB code (R2024b, MathWorks, Inc.) and code developed for the EEGLAB toolbox^81^ and the Brainstorm toolbox^82^.

#### EEG power spectrum density analysis and FOOOF analysis

We computed the power spectrum density (PSD) for the alpha (8–13 Hz) and beta (14-30 Hz) band using the modified Welch periodogram method implemented in the MATLAB ‘pwelch’ function and a fast Fourier transform (FFT). To have a nominal frequency of 0.125 Hz, we segmented the preprocessed EEG signal into 6-s windows with zero padding and 50% overlap. Next, we converted the absolute PSD values into relative power by dividing each participant’s alpha power by the total power across all frequencies included in the analysis, allowing for direct comparisons between subjects and conditions. We segmented the EEG signal in 1000-ms epochs (from -500 to +500 relative to each touch applied to the participants’ hand). Power was normalized by z score decibel transformation. We selected electrodes over the somatosensory cortex, the posterior parietal cortex and the premotor cortex. Indeed, beta oscillations in the somatosensory and parietal cortices are reliably observed in response to tactile stimulation^45–47,52,53,62–64^. Moreover, the posterior parietal cortex and premotor cortex play a key role the tactile integration underlying body ownership^30^. For the somatosensory and posterior parietal cortex, we defined one parietal region of interest (ROIs) covering both hemispheres: D14; D15; D16; A6; A5; A7; A18; A8; A17; A9; A16; B1; B20; B2; A32; B3; A31; B4; A30; B5; A29; B6. In addition, we selected a premotor ROI corresponding to electrodes over the bilateral premotor cortex, including its ventral and dorsal parts: C26; C25; C24; D2; C32; D4; D3; D13; D7; D6; D5; D12; D11; D10; D9; D8; C13; C12; C11; C2; C10; C4; C3; B32; C7; C6; C5; B31; B30; B29; B28; B27. These data were subsequently exported for statistical analysis. PSD and FFT were performed using custom-made MATLAB code and code developed for the Brainstorm toolbox^63^. We implemented the recent “Fitting Oscillations and One Over F” (FOOOF) algorithm to increase the precision of periodic estimates by modeling brain oscillations after excluding the aperiodic component^54,55^. This allowed us to increase the precision of the oscillation estimates by excluding the aperiodic component of the EEG spectrum. We applied the FOOOF algorithm^54^ to Welch’s power spectral density (PSD) data using the FOOOF/Specparam function in Brainstorm for MATLAB^81^. This algorithm was implemented separately on each electrode for the alpha (8–13 Hz) and beta (14-30 Hz) frequency range with a sliding time window with 50% overlap to fit spectral peaks, using default parameters: a Gaussian distribution, a minimum number of peaks of 3, a minimum peak height of 1 dB, and a proximity threshold of two standard deviations from the largest peak. To assess the FOOOF model’s performance, we quantified the goodness of fit (R²) for the body ownership, simultaneity judgment, and resting-state conditions across all electrodes of interest. We then extracted peak parameters from the periodic component and calculated the IBF and IAF.

The FOOOF/Specparam algorithm successfully identified a beta peak (14–30 Hz) in most of the 22 electrodes within the parietal ROI across all participants in the resting-state condition (mean number of electrodes with a peak: 21.58; range: 17–22), during body ownership judgments (mean: 21.31; range: 14–22), and during simultaneity judgments (mean: 21.42; range: 17–22). Similarly, FOOOF detected alpha peaks (8-13 Hz) in most of the 22 electrodes comprising parietal ROI during the resting-state condition (mean: 21.92; range: 20–22), as well as during both body ownership and simultaneity judgments, where a peak was identified in all electrodes. For the premotor ROI, the FOOOF/Specparam algorithm successfully identified beta peaks for the majority of the 32 electrodes across all participants in the resting state condition (mean: 31,138 range: 22-32), in the body ownership judgments (mean: 30,333; range: 25-32) and in the simultaneity judgments (mean: 30,694; range: 25-32). Similarly, the FOOOF model detected alpha peaks in most of the electrodes during the resting-state condition (mean: 31,833; range: 27–32), as well as during body ownership (mean: 31,833; range: 27-32) and during simultaneity judgments (mean: 31,777; range: 24-32).

#### Data analysis: assessing correlations of IBF and IAF with TBWs and sensitivities

We ascertained correlations by computing Pearson and Spearman correlations between TBWs between IBF or IAF measured in our selected ROIs during both resting-state recordings and perceptual judgment tasks. We additionally examined correlations between IBF or IAF and perceptual sensitivity for body ownership and tactile simultaneity. To this end, we employed the same correlational approach for the sensitivity to body ownership and simultaneity judgment tasks (AUC d’).

### Experiment 3

#### Participants

Participant sample consisted of the same individuals (N = 46; 26 women; age range 18–43 years; mean age = 27.98 years) tested in our previous study^27^. The original dataset was reanalysed here to investigate whether IBF predicted body ownership and simultaneity TBWs.

#### Experimental setup and procedure

The experiment was conducted in the same room dedicated to EEG recording as Experiment 2. Prior to performing the perceptual tasks in the main experiment, we recorded eight minutes of resting-state EEG: four minutes with eyes closed and four minutes with eyes open. For the subsequent analyses, we considered only the eyes-closed resting-state condition to mirror the analyses performed in Experiment 2. Like Experiment 1 and 2, participants engaged in two different tasks: *(i)* a body ownership judgment task and *(ii)* a simultaneity judgment task. The order of the tasks was counterbalanced across participants. In both tasks, participants maintained their right-hand out of view, lying palm down on a flat support 30 cm lateral to the body midline. A chin rest and an elbow rest ensured that the participants’ head and arm remained in a steady and relaxed position. During body ownership judgment tasks, a rubber right hand, a cosmetic prosthetic right hand (model 30916-R, Fillauer®) filled with plaster, was placed in view on a platform 15 cm directly above and aligned with the real hand that was placed on a lower platform out of view. Two custom-made robotic arms applied tactile stimuli (taps) to the rubber hand’s index finger and to the participant’s hidden real index finger (for details^9,12^). Taps on the rubber hand were either synchronized with the taps on the participant’s real hand (synchronous condition) or were delayed or advanced at different asynchronies. The asynchronies varied across trials in five steps between ±400, ±300, ±200, ±100, and 0 (true synchrony) ms, where a negative value indicates that the rubber hand was touched first, and a positive value indicates that the participant’s hand was touched first. During stimulation participants wore earplugs to minimize distraction from any noise produced by the robots and they were instructed to focus their gaze on the rubber hand.

In each trial, the robotic arms tapped the rubber hand’s index finger and the participant’s index finger six times each for a total period of 12 s. The robotic arms tapped the corresponding parts of the real and rubber fingers, targeting four different locations in randomized order (immediately proximal to the nail on the distal phalanx, on the middle phalanx, on the proximal interphalangeal joint, and on the proximal phalanx). The interval between one touch and the other was jittered between 1100 and 1300 ms to avoid that a regular rhythm might cause cortical oscillation entrainment. Then, the robots stopped while the participant heard a tone instructing them to verbally provide their answer. In the body ownership judgments participants reported whether the rubber hand felt like their own hand by saying “yes” (“the rubber hand felt like it was my hand”) or “no” (“the rubber hand did not feel like it was my hand”). In the simultaneity judgments participants reported if the touches on their real hand and the touched on the rubber hand were synchronized or not. Five seconds later, a second tone informed the participant that the next trial was about to start, and the next trial started 1 s after this sound cue. As in Experiment 2, body ownership and simultaneity judgment tasks were administered in three different blocks of trials using the ABAABB counterbalancing order, with the order counterbalanced across participants. Each asynchrony was repeated 5 times in each block, leading to a total of 45 trials per block. In total, each asynchrony was repeated 15 times in both the body ownership and the simultaneity judgments.

#### EEG recording, preprocessing, power spectrum density and FOOOF analysis

EEG recording and preprocessing procedures were identical to those described for Experiment 2, as was the power spectral density analysis. As in Experiment 2, we segmented the EEG signal into 1000-ms epochs (from –500 to +500 ms relative to each touch applied to the participant’s hand) and considered the same ROIs used in Experiment 2. FOOOF parameters were also identical to those in Experiment 2, with the only difference being that we extracted only IBF (14-29 Hz), as a correlation between IAF and visuotactile integration had already been demonstrated in^12^. Also in this case, the FOOOF/Specparam algorithm successfully identified a beta peak (14–30 Hz) in most of the 22 electrodes within the parietal ROI across all participants in the resting-state condition (mean number of electrodes with a peak: 21.87; range: 17–22), during body ownership judgments (mean: 19.93; range: 10–22), and during simultaneity judgments (mean: 19.00; range: 8–22). For the premotor ROI, the FOOOF/Specparam method successfully identified beta peaks for the majority of the 32 electrodes across all participants in the resting state condition (mean: 31,804 range: 25-32), in the body ownership judgments (mean: 30,543; range: 23-32) and in the simultaneity judgments (mean: 29,500; range: 13-32).

#### Data analysis: assessing correlations of IBF with TBWs and sensitivities

We ascertained correlations by computing Pearson and Spearman correlations between TBWs and the IBF measured in our selected ROIs during perceptual judgment tasks and during the resting-state period. We additionally investigated possible correlations between IBF and sensitivities to body ownership and visuotactile simultaneity. Therefore, we employed a similar correlational approach for the sensitivity to body ownership and simultaneity judgment tasks (AUC d’).

## Acknowledgments

We would like to thank Martti Mercurio for writing the program to control the robots, Mattias Karlén for creating the illustrations used in Figures 1 and 4, and Michelangelo Tani for his help in collecting the data of Experiment 1. H.H.E. was supported by a Swedish Research Council Distinguished Professor grant (VR; #2017-03135), Hjärnfonden and Torsten Söderberg’s stiftelse. M.D. was supported by H2021 Marie Skłodowska-Curie Actions (Grant agreement no 101063812) and by Sweden’s Innovation Agency (VINNOVA; #2022-01441). R.C.L. was supported by a Swedish Research Council (VR; #2024-00839) project grant and by a Strategic Research Area Neuroscience (StratNeuro) research fellowship.

## Author contributions

M.D. and H.H.E. conceived and designed the study. M.D. collected the data, analyzed the data and created the figures. R.C.L. created the code for EEG data analysis and analyzed EEG data. M.D. and H.H.E. wrote the manuscript. All authors edited the manuscript.

