## Supplementary Material for "Beta oscillations support somatosensory temporal integration for body ownership"

**Supplementary Methods**

**Participants inclusion test**

Participants who did not experience a clear and reliable somatic rubber hand illusion were unable to make reliable body ownership judgments. Therefore, all participants underwent an inclusion test to ensure they were capable of experiencing the illusion and could meaningfully perform the body ownership judgments**.** In the inclusion test, we used the same haptic robot devices used in the main experiment. As in the main experiment, participants’ left index finger was attached with a velcro strap to a custom-made support attached to the stylus of one haptic device, which guided their finger to perform tapping movements on the silicone hand also covered with a white glove. A second haptic device held a fake wooden finger, likewise covered with a piece of the same white glove and fitted with a silicone layer beneath the glove at the contact site with the participant’s real hand. This device applied taps to the participant’s real right hand, specifically on the dorsum of the hand, between the third and fourth metacarpal bones, in an area free of bony prominences to ensure uniform tactile stimulation. In the inclusion test, the stimulation consisted of a synchronous trial comprising12 touches over 24 seconds. At the end of a single 24 s stimulation, participants were asked to complete a 5-items questionnaire to assess the feeling of ownership (**Table S1**). Participants were asked to indicate the extent of their agreement or disagreement with six statements using a seven-point Likert scale, ranging from −3 (“I completely disagree”) to +3 (“I completely agree”), with a response of 0 indicating “neither agreed nor disagreed”. One statement examined the perception of the illusion (Q1) and the remaining four statements were designed to control suggestibility and task compliance (Q2-Q5). Our inclusion criteria for a somatic rubber hand illusion strong enough for participation in the main psychophysics experiment were as follows: (i) a score for the illusion statement greater than 1 and (ii) a difference between the score for the illusion item and the mean score for the control items greater than 1.

| Q1 | It felt like I was touching my right hand with my left index finger |
| --- | --- |
| Q2 | It felt like I had more than one right hand |
| Q3 | It felt like my right hand was larger than normal |
| Q4 | It felt like my right hand was moving |
| Q5 | It felt like I was not able to feel my own right hand |

**Table S1.** **Questionnaire statements used in the inclusion test**. Q1 assesses the specific illusory experience of interest (i.e., the sensation of touching one’s own hand). The remaining statements (Q2–Q5) serve as control items to assess suggestibility and task compliance
